# Virus-Like Particles: The Next Frontier in Livestock Gene Editing

**DOI:** 10.64898/2026.03.30.715406

**Authors:** Theresa M. Pauli, Theresa von Heyl, Beate Rieblinger, Sabrina Teresa Schleibinger, Wei Liang, Antonia Schmauser, Mithuoshni Arullmoli, Paul Derrer, Anika Eckstein, Sirradu Jagana, Camila Gatti Corrêa, Krzysztof Flisikowski, Tatiana Flisikowska, Benjamin Schusser

## Abstract

Pigs and chickens are not only the most important livestock species for global food production but also serve as key model organisms in various research disciplines. The pig is widely used in translational research due to its anatomical and physiological similarity to humans, providing valuable insights into immunology, metabolism, and disease mechanisms. In contrast, the chicken has become an essential model for studies related to poultry health, animal welfare, and developmental biology. Its externally developing embryo offers exceptional accessibility for experimental manipulation. Recent advances in genome editing technologies, particularly CRISPR/Cas9, have further expanded the potential of these species for functional genomic studies, although the efficient delivery of such tools remains a major challenge. By using virus-like particles (VLPs), we have been able to overcome this limitation. Here, we evaluated VLPs as delivery vehicles for genome engineering tools in pigs and chickens, two key livestock species at the human–animal interface. VLP-mediated delivery enabled efficient Cre recombination and high CRISPR/Cas9 editing rates in porcine cells, organoids, and oocytes, particularly when multiplexed. In chickens, VLPs supported robust Cre recombination and Cas9-mediated editing in cell culture, tracheal organ cultures, and *in ovo*. Reporter VLPs and dCas9 VLPs further demonstrated the versatility of this platform across porcine and avian systems. Together, these findings establish VLPs as an efficient and time-saving strategy for gene editing in livestock, with relevance for animal health, agricultural productivity, and translational One Health research.

## Introduction

Chickens and pigs are not only essential sources of animal-derived protein in human nutrition but also valuable models in research fields such as immunology, infectious diseases, cancer, and developmental biology. In recent years, pigs have emerged as the most important translational model organism, bridging preclinical research and human trials (1). They are used in various biomedical research fields such as diabetes (2), cancer (3), or cognitive and behavioral research (4). Compared to rodents, they offer many significant advantages. Of particular importance are their relatively long lifespan, which enables longitudinal studies, as well as the closer resemblance to the human pathophenotype (5). The similarity in size, physiology, and anatomy supports the advancement of surgical research and additionally attributes great translational value to the pig (6). This also includes the field of xenotransplantation, driven by the critical shortage of human donor organs, for which the pig is considered the most suitable donor species (7). As a result, the demand for genetically modified (GM) pigs has increased significantly over the past decade. Similarly, chickens have gained substantial importance as complementary model organisms in both basic and applied research. GM chickens play a central role in developmental biology (8), where they have enabled fundamental discoveries in embryogenesis and tissue differentiation. They have provided key insights into immunology (9, 10) and mechanisms of disease resistance (11, 12), contributing to both veterinary and human health research. Compared to mammalian models, chickens offer unique experimental advantages, including easy access to the developing embryo and cost-effective large-scale studies. In addition, they serve as efficient bioreactors for the production of recombinant proteins *in ovo* (13), further increasing their biotechnological and translational value. Taken together, valuable models for biomedical and translational research have been established using chickens and pigs. Nevertheless, the efficiency and safety of generating such models could be greatly improved by advancing delivery tools for these species (14).

CRISPR/Cas9 and Cre recombinase are widely used genome engineering tools that have been substantially refined in recent years. Innovations such as catalytically inactive Cas9 (dCas9), base editors, and transcriptional modulators (CRISPRa/i) have expanded the range of possible genetic modifications beyond simple gene disruption (15). These technologies now form the foundation of state-of-the-art strategies for generating GM animals. In pigs, the historical lack of fully functional germline-competent embryonic stem (ES) cells necessitated gene targeting in somatic cells, a technically demanding and inefficient approach (16). This limitation significantly restricted the generation of GM pigs until the introduction of CRISPR/Cas9, which enabled direct and more efficient genome editing. Today, GM pigs are typically produced either by microinjection of genome-editing tools into zygotes or by modifying somatic cells followed by somatic cell nuclear transfer (SCNT) (17). A comparable challenge existed in chickens, where ES cells also failed to contribute to the germline. This obstacle was overcome through the possibility to cultivate and manipulate germline-competent primordial germ cells (PGCs), thereby allowing precise genome engineering, including targeted knock-ins and knock-outs (9, 18). Currently, genetic modification in chickens relies predominantly on the editing of PGCs (19). Although these approaches are well established, they remain labor-intensive and time-consuming, particularly due to the need to breed founder animals, and they may result in unintended genotypes (14). To better comply with the 3R principles and accelerate research timelines, there is a growing demand for efficient, safe, and rapid *in vivo* delivery systems for genome engineering tools (20). These strategies would allow more precise and time-efficient generation of modified animals, as they enable the refinement of existing animal models while reducing the need to generate new ones.

Virus-like particles (VLPs) are self-assembling nanostructures ranging from 20 to 200 nm in size, with broad applications in therapeutics, vaccines, and diagnostics (21). In mouse models, VLPs have been successfully used as protein delivery vehicles for genome engineering tools such as CRISPR/Cas9 and Cre recombinase not only *in vitro* but also *in vivo* (22, 23). Importantly, VLP-based delivery has proven particularly effective for the *in vivo* editing of immune cells, demonstrating the feasibility of targeting key components of the immune system with high efficiency (24). While this delivery system is well established in mice, there is no evidence of the effectiveness of genome engineering using VLPs in livestock species such as pigs and chickens. In this study, we systematically evaluated a repertoire of these nanostructures including Cre and Cas9 VLPs in these species. We demonstrated that in both pigs and chickens, VLPs provide an efficient platform to deliver genome editing tools and fluorescent proteins *in vitro* in cells and organoid systems, *ex vivo* in organ cultures and additionally *in ovo* in chicken. The use of this technology will advance the development of genetically modified livestock models, contributing to disease prevention and improved animal health according to the One Health approach.

## Results

### Efficient VLP-mediated delivery of fluorescent proteins into somatic and primordial germ cells

The first goal was to test the transduction efficiency of VLPs on porcine primary kidney fibroblasts (pKDNFs), chicken DT40 cells, as well as chicken primordial germ cells (PGCs) by delivery of fluorescent proteins. pKDNFs, DT40 cells and PGCs were transduced with VLPs packaged with superfolder (sf) GFP. As early as 24 h post transduction, FACS analysis and fluorescence microscopy showed that nearly all somatic porcine and chicken cells (>90%) were successfully transduced (Figure 1A and B). After 48 h, only a slight decrease in the fluorescence of sfGFP signal in pKDNFs was visible. Additionally, VLPs carrying the red fluorescent protein mCherry transduced pKDNFs and DT40 cells with efficiencies comparable to those of sfGFP VLPs (supplementary figure S1A and C). Given that PGCs are known to be difficult to transfect, varying VLP volumes were evaluated in PGCs. It was found that 20 µL VLPs reached comparable results to 5 µL VLPs on somatic cells (Figure 1A, supplementary figure S1B and D).

**Figure 1.**
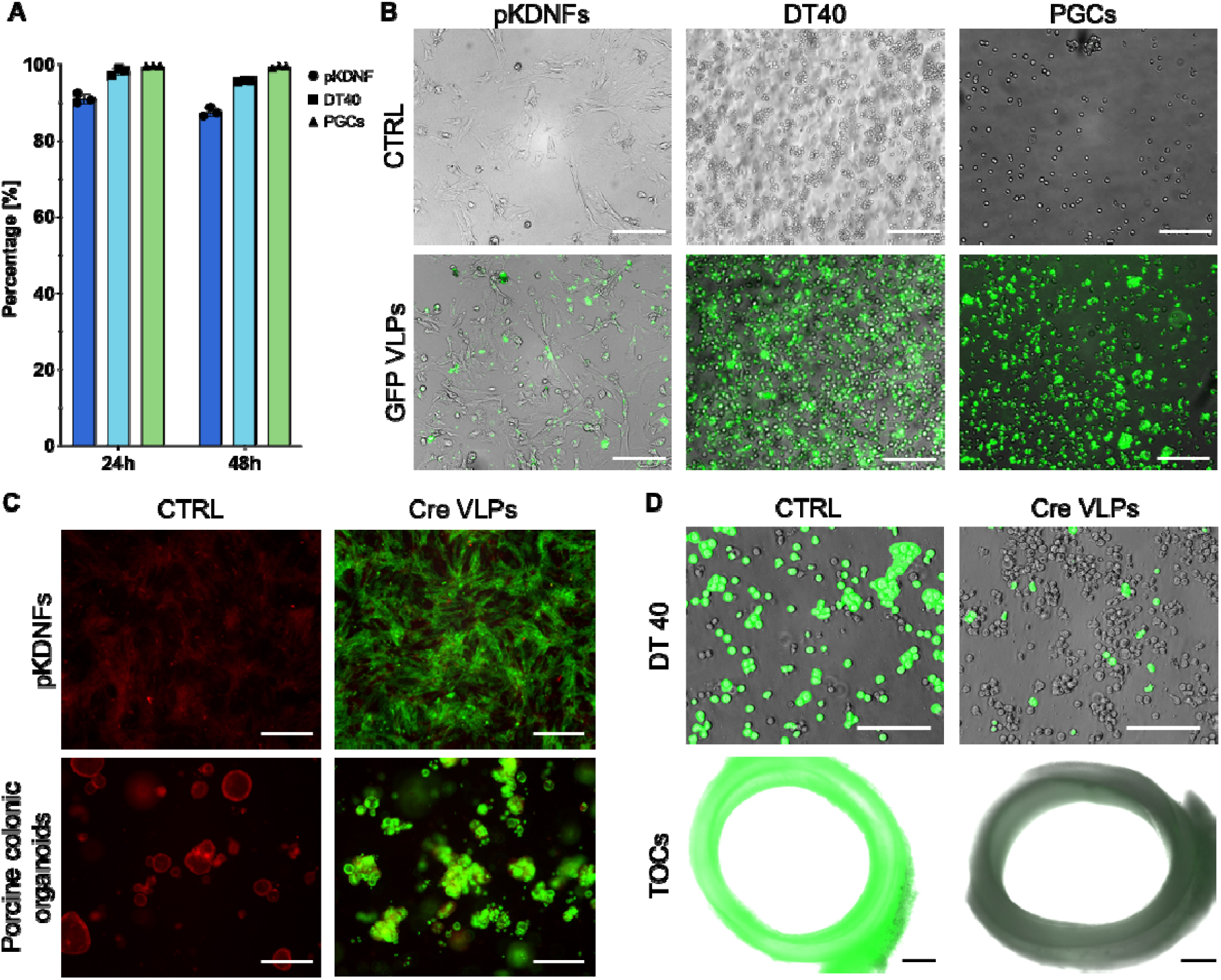
VLPs efficiently package and transport fluorescence proteins and Cre reombinase and release them into various target cells *in vitro* and *ex vivo*. **A:** Relative count of fluorescence expression in single cells from porcine pKDNFs and DT40 chicken cells transduced with 5 µL sfGFP VLPs and chicken PGCs transduced with 20 µL sfGFP VLPs 24 h and 48 h post transduction compared to control (n=3; SD shown). **B**: Representative fluorescence microscopy images of pKDNF and DT40 cells transduced with sfGFP VLPs (5 µL) and chicken PGCs transduced with 20 µL sfGFP-VLPs 24 h post transduction. Scale bar = 130 µm. **C**: Representative fluorescence microscopy images at 48 h post transduction of pKDNFs (top) and porcine colonic organoids (bottom) from a dual-fluorescent reporter pig. Control (CTRL) samples show constitutive mTomato expression, and eGFP expression occurs after successful delivery of Cre recombinase via VLPs. 10 µL (pKDNFs) or 5 µL (organoids) of Cre VLPs were used. Scale bar = 420 µm. **D**: Representative fluorescence microscopy images of *Igh*^KO^ DT40 chicken cells (top) transduced with 20 µL Cre VLPs 24 h post transduction. Scale bar = 230 µm. Representative fluorescence microscopy images of TOCs (n=3) from JH-KO chicken embryos (bottom) 48 h post transduction with 10 µL Cre VLPs. Scale bar = 430 µm.

### VLP-delivered Cre efficiently excises loxP flanked regions in porcine and avian cultures

To test the functionality of Cre VLPs on pKDNFs, we used cells from a dual-fluorescent reporter pig (25), where successful recombination excised a loxP-flanked stop (*LSL*) cassette containing the *mTomato* gene, and thereby activating downstream eGFP expression as an easy and direct readout. Cre VLPs produced strong expression of eGFP 48 h post transduction (Figure 1C top), indicating successful excision of the *LSL* cassette. We extended this study to evaluate VLPs in a non-mammalian vertebrate, the chicken. To assess the efficiency of Cre VLPs in chicken, *Igh*^KO^ DT40 cells – a chicken B cell lymphoma line in which the immunoglobulin heavy chain (*Igh*) gene is disrupted (26) – were used. Transduction of these cells with Cre VLPs was expected to result in recombination of the *Igh*^KO^ repair template cassette and simultaneous excision of eGFP. Accordingly, fluorescence microscopy revealed that Cre VLPs resulted in only a minor detectable eGFP signal in any transduced group after 24 h, which indicated efficient recombination and consequently, loss of eGFP expression (Figure 1D top; supplementary figure S2A). Flow cytometry analysis demonstrated that 24 h post transduction, a significant decrease in eGFP-positive cells was observed in the group transduced with 20 µL Cre VLPs. Similarly, after 72 h a significant decrease was also observed in the group transduced with 10 µL compared with the control (supplementary figure S2B).

To test Cre VLPs on a more complex *in vitro* system, we investigated their functionality on porcine colonic organoids from the same reporter pig. Transduction protocols for human and murine organoids (27) were adapted and optimized for porcine organoids leading to efficient recombination caused by Cre VLPs. Strong eGFP expression was evident after 48 h (Figure 1C bottom). As a more elaborate avian *ex vivo* culture system, tracheal organ cultures (TOCs) from JH-KO chickens (9) were used. Similar to the principle described above, VLP-mediated delivery of Cre led to recombination and consequent loss of eGFP signal. A decrease of eGFP fluorescence in the VLP treated TOCs was visible 48 h after transduction (Figure 1D bottom).

To assess whether the baboon envelope (BaEV) glycoprotein enhances transduction of porcine cells as it is the case for other species (22), Cre VLPs were produced with varying ratios (3:0, 2:1, 1:2, 0:3) of vesicular stomatitis virus glycoprotein (VSV-G) and BaEV. The distinctly pseudotyped VLPs were used to transduce pKDNFs and in the recommended ratio of 1:2 VSV-G:BaEV, the transduction efficiency was strongly decreased and resulted in less eGFP signal while the ratio of 3:0 VSV-G:BaEV showed the best result (supplementary figure S3). Consequently, all experiments in this study were conducted using VSV-G only.

To quantify protein content in the VLPs and determine the optimal amount of VLPs for transduction, three distinct batches of Cre VLPs were analyzed through ELISA and additional functional tests on pKDNFs (supplementary figure S4). Based on a positive correlation, 5 -10 µL of Cre VLPs with a concentration of 2.55 U/µL were used in the experiments.

### Efficient Cre VLP–mediated activation of latent *KRAS* and *TP53* mutations in porcine organoids

After Cre VLP functionality was proved in a fluorescent reporter system, organoids from pigs that carry latent mutations in the oncogene *KRAS* and the tumor suppressor gene (TSG) *TP53* (28, 29) were established. The oncogenic mutations *KRAS*^*G12D*^ and *TP53*^*R167H*^ are silenced by a *LSL* cassette in the first intron of the gene. Organoids were transduced with Cre VLPs and grown in culture for up to 46 days. From day 24 in culture, we observed a change in the organoids’ morphology from a branched, differentiated to a cystic and proliferative architecture (Figure 2A, supplementary Figure S5 and S6) indicating activation of the silenced mutations. Recombination PCR was performed to assess excision of the *LSL* cassette in both genes. As early as day 5, recombined *TP53*^*R167H*^ (Figure 2B) and *KRAS*^*G12D*^ (Figure 2C) alleles were detectable, with increasing intensity over time. By day 26, the *LSL* cassette was no longer amplifiable, indicating complete excision. For *KRAS*, loss of the WT band suggested loss of heterozygosity at DNA level. To evaluate transcriptional consequences, RNA was converted to cDNA and allelic expression was quantified. For both *TP53* (supplementary figure S7A) and *KRAS* (supplementary figure S7B), WT and mutant transcripts were still detectable at later time points (day 20 and day 26, respectively). Together, these data demonstrate that VLP-mediated Cre delivery enables stable activation of latent oncogenic alleles in porcine organoids and drives their phenotypic transformation.

**Figure 2.**
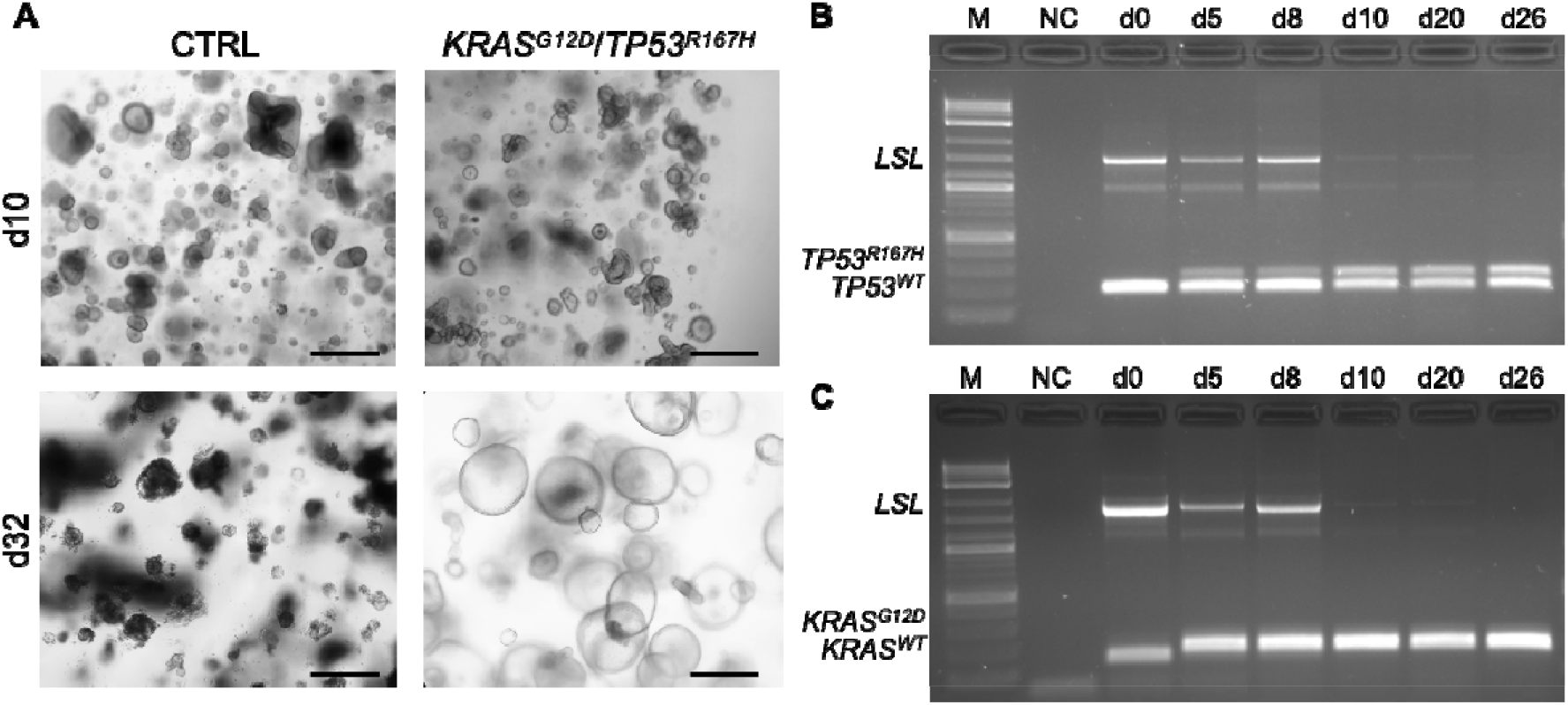
VLPs efficiently deliver functional Cre to porcine colonic organoids to activate latent oncogenic mutations. **A**: Representative brightfield images of organoids that were transduced with 5 µL Cre VLPs to activate latent oncogenic mutations in *KRAS* and *TP53* at d10 and d32 post transduction. At d10, organoids of CTRL and VLP treated groups display a similar (early) branched, differentiated structure. At d32, unedited organoids are entering the burned-out stage, wheras edited organoids change their morphology to a cystic, proliferative architecture. Scale bar = 420 µm. **B** and **C**: Products of *TP53* (**B**) and *KRAS* (**C**) recombination PCR pre (d0) and post Cre recombination visualized on agarose gels. Representative results of organoid samples from a pig with latent mutations in *TP53* and *KRAS*. After Cre VLP transduction, the *LSL* cassette is excised and a band representing the activated mutant allele appears. Mutant band intensity increases until the band for the *LSL* cassette disappears completely. M: marker, NC: negative control.

### Multiplexing strategy leads to high genome editing in porcine cells, organoids and oocytes

As a next step toward genome editing in livestock, Cas9 VLPs were applied to pKDNFs targeting different TSGs to generate single knockouts (SKOs). 48 h after transduction, inference of CRISPR edits (ICE) analysis of isolated gDNA revealed high editing efficiencies, with insertion and deletion (INDEL) rates of 80–98% and corresponding knockout (KO) scores of 70–98% (Figure 3A). To achieve even higher KO scores more consistently, different multiplexing strategies were evaluated (supplementary figure S8A). Inclusion of two gRNAs in a single VLP batch (multiplexing strategy 1, MPS1) markedly reduced editing efficiency and showed a bias towards one gRNA (supplementary figure S8B). In contrast, co-transduction with two separately produced VLP batches, each containing one gRNA (MPS2), resulted in editing efficiencies comparable to SKOs and was therefore used for subsequent experiments (supplementary figure S8B). Using MPS2, dual targeting of *IL-10* with flanking gRNAs achieved INDEL and KO scores of up to 100% in pKDNFs, with ICE analysis confirming complete excision of the target region (supplementary figure S8C, D, and E). MPS2 was further applied to porcine organoids, multiplexing Cas9 and Cre VLPs simultaneously. Five days after transduction, ICE analysis showed an INDEL score of 81% for *B2M*, comparable to a SKO, while simultaneous Cre activity and successful recombination was confirmed by strong eGFP expression (supplementary figure S8C, D, F, and G). Finally, multiplexed Cas9 VLPs targeting *IL-10* were microinjected into *in vitro* fertilized porcine oocytes (supplementary figure S9A and B). After 7 days in culture, 66.6% of blastocysts were edited, with INDEL rates of 94–100% and KO scores of 61–100% (Figure 3B). In one blastocyst, 100% KO was achieved, with 98% of edits resulting from dual gRNA-mediated excision, as confirmed by ICE analysis (Figure 3C).

**Figure 3.**
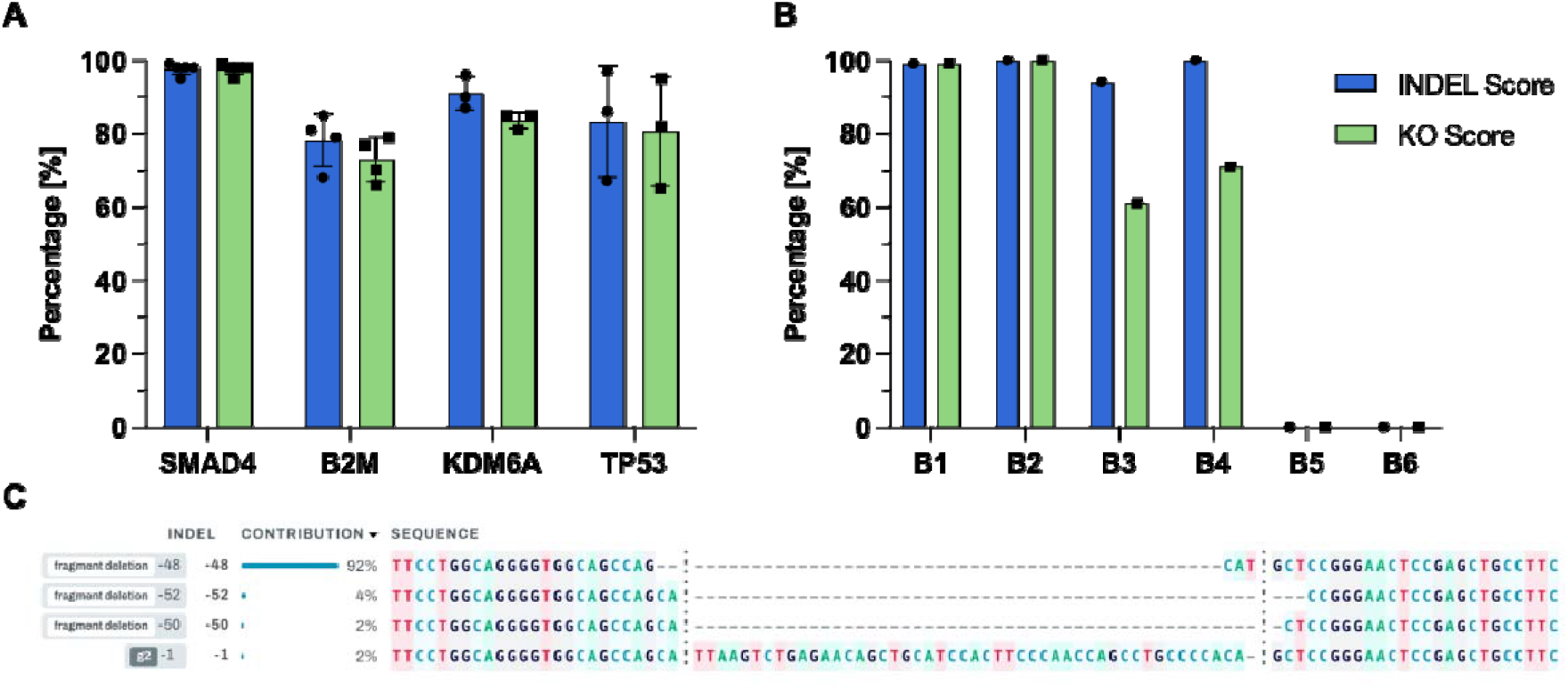
Cas9 VLPs efficiently generate KOs in porcine fibroblasts and blastocysts. **A**: Bar graph showing ICE analysis results of targeting of *SMAD4, B2M, KDM6A* and *TP53* in pKDNFs using Cas9 VLPs. Blue bars: INDEL scores, green bars: KO scores compared to WT control (n≥3; SD shown). **B**: Bar graph showing ICE analysis results of targeting of *IL-10* in developed blastocysts (B1-B6) using multiplexed Cas9 VLPs. Blue bars: INDEL scores, green bars: KO scores compared to WT control. **C**: Detected INDELs of blastocyst 2 (B2) report that in 98% of the cases, the whole fragment was deleted.

### *In vitro* and *in ovo* delivery of Cas9 VLPs directed against chicken *B2M* successfully reduces MHC-l expression

To evaluate Cas9-mediated genome editing in chicken cells via VLPs, the *B2M* locus was targeted. MHC-I surface expression on adherent HD11 cells and in the bursa of embryonic day (ED)19 embryos was analyzed by flow cytometry following transduction with Cas9 VLPs. A pronounced reduction in mean fluorescence intensity (MFI) was observed in HD11 cells at 48 h and 72 h post transduction (Figure 4A). Furthermore, gene editing in chicken PGCs achieved a mean INDEL frequency of 20% at 72 h post transduction (Figure 4D; supplementary figure S10B). For *in ovo* delivery, *B2M*-targeting VLPs were injected intravenously into ED12 chicken embryos (Figure 4C). Gene editing was assessed seven days later (ED19). A significant decrease in surface MHC-I expression was detected in the bursa (Figure 4B), whereas reduced MHC-I expression in PBMCs was observed in a subset of embryos (supplementary figure S10A), indicating tissue-specific editing efficiencies. Notably, all analyzed bursi (9/9; 100%) exhibited detectable gene editing, with 7/9 reaching INDEL frequencies >10% (supplementary figure S10C), underscoring the robustness of the *in ovo* VLP delivery.

**Figure 4.**
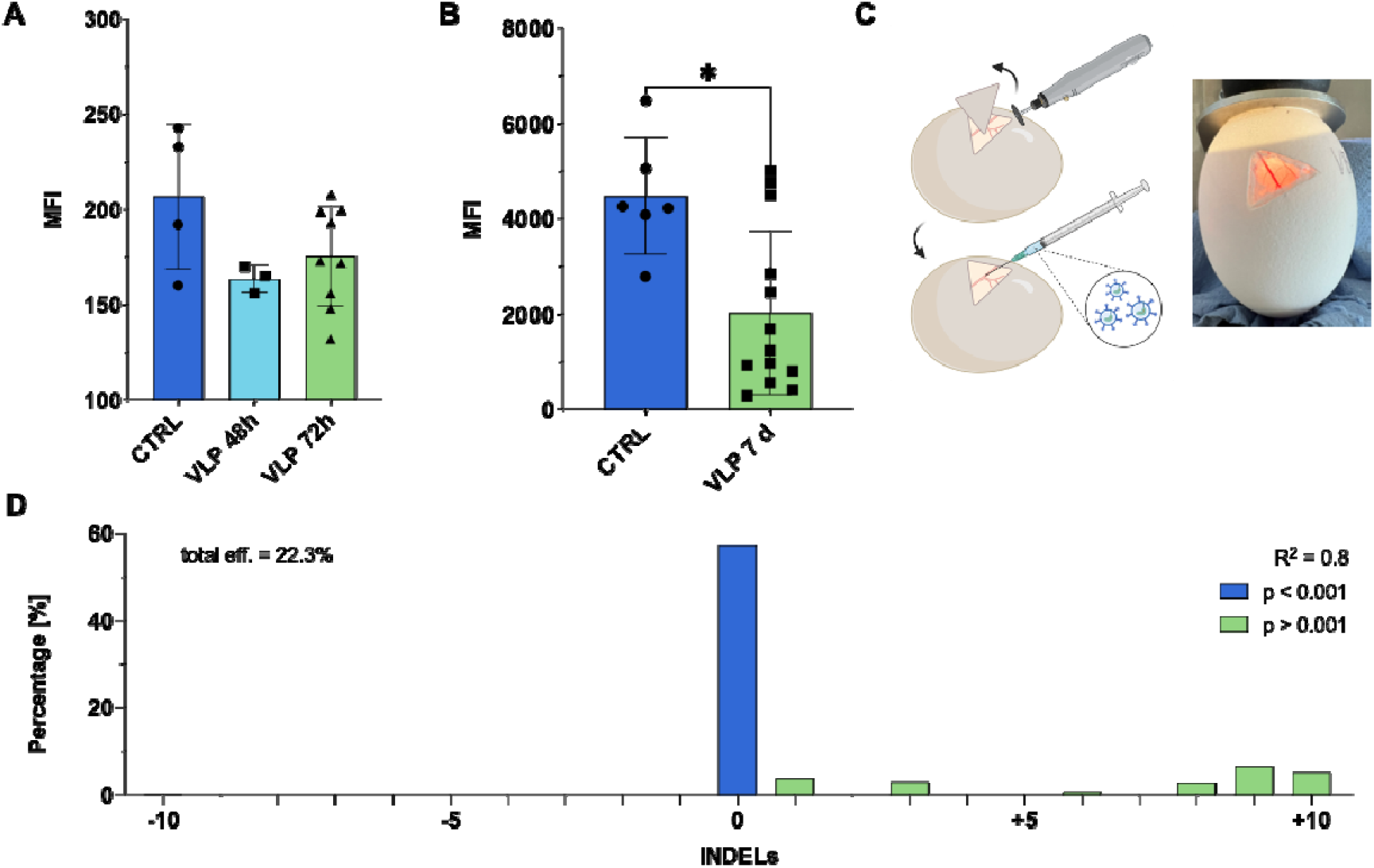
VLP-mediated gene editing decreases MHC-l expression on the surface of chicken cells *in vitro* and *in ovo*. **A**: Mean fluorescence intensity (MFI) of MHC-I expression in HD11 chicken cells transduced with 10 µL Cas9 VLPs targeting *B2M* at 48 h and 72 h post transduction compared to control. (n≥3; p<0,05; Multiple ANOVA; SD shown). **B**: MFI of MHC-I expression in bursocytes from ED19 WT chicken embryos transduced with 50 µL Cas9 VLPs targeting *B2M* 7 days post transduction compared to uninjected control (n≥6; p<0,05; Multiple ANOVA; SD shown). **C**: Schematic workflow of *in ovo* injection of Cas9 VLPs. Briefly, a triangle is cut into the eggshell without destroying the membrane; 50 µL VLPs are then injected into a vein using an insulin 30 G syringe. **D**: Representative TIDE analysis of cells from PGCs transduced with Cas9 VLPs targeting *B2M* 72 h post transduction.

### Editing the epigenome using dCas9 VLPs in porcine cells

To enable epigenome editing, a plasmid for expression of a Gag-dCas9 fusion protein was cloned using the plasmid for Gag-Cas9 expression as template for site-directed mutagenesis. dCas9 VLPs were used to test a passive demethylation approach proposed by Sapozhnikov et al. (30). Therefore, intron four in *TP53* was targeted, since a highly methylated internal promoter (P2) is located in this region. Two established gRNAs were used to target several CpG islands of interest (Figure 5A). Multiplexed dCas9 VLPs led to a demethylation frequency of up to 21% in pKDNFs after 6 days (Figure 5B). CpGs 2 and 3 showed the highest demethylation rate because they are targeted by both gRNAs simultaneously, and the results align with the predicted demethylation window. The use of dCas9 VLPs including only gRNA1 led to less demethylation, even after a prolonged transduction period of 15 days (supplementary figure S11).

**Figure 5.**
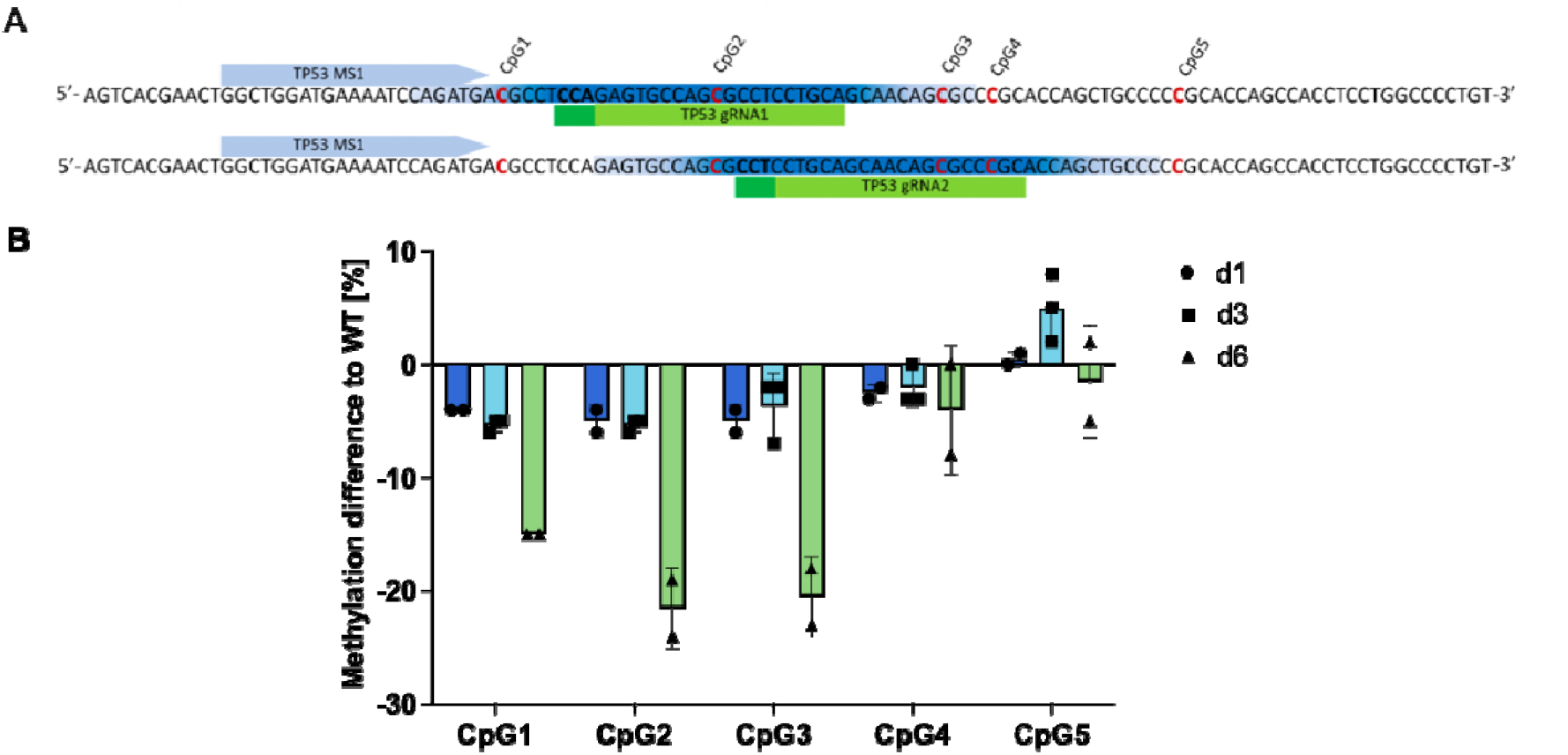
Epigenome editing using multiplexed dCas9 VLPs targeting internal P2 in *TP53*. **A**: The internal promoter P2 of *TP53* is targeted using two gRNAs that bind to regions with highly methylated CpG islands, leading to dCas9 blocking the locus for further methylation. Light green: gRNA, dark green: PAM, light blue: sequencing primer, dark blue: expected demethylation window, red: CpG islands of interest. **B**: pKDNFs were transduced every second day with multiplexed dCas9 VLPs and analyzed on day 1, day 3, and day 6 post transduction. Methylation was measured compared to WT in percentage (n=2-3; SD shown).

## Discussion

In this study, we establish VLP-mediated delivery as an efficient platform for transferring diverse cargoes into cells from multiple livestock species. Through optimization of VLP production, we generated a versatile panel that performed robustly across *in vitro, ex vivo*, and *in ovo* systems. This approach addresses key barriers that have limited the translation of genome engineering technologies from model organisms to livestock, with important implications for model refinement, scalability, and ethically responsible research.

Efficient VLP production for livestock applications required systematic optimization of Gag and VSV-G expression in producer cells compared to murine and human systems (22, 27). The recruitment of envelope glycoproteins such as VSV-G by Gag during virion formation is a complex process that remains incompletely understood. Current evidence indicates that no single Gag domain mediates this process, but rather that VSV-G and other envelope proteins are recruited at lipid-rich assembly and budding sites (31). Higher levels of VSV-G expression in producer cells increase its incorporation into virions and can enhance transduction efficiency by up to 55-fold *in vitro* (32). Implementing these adjustments enabled robust transduction of porcine and chicken cells, organoids, organ cultures, and oocytes, underscoring the need for species-specific optimization when adapting VLP technologies to livestock.

Reporter VLPs carrying sfGFP or mCherry efficiently transduced porcine and avian somatic cells as well as PGCs without detectable toxicity. In addition to serving as process controls, these reporters enabled cell sorting and rapid verification of successful delivery, particularly in PGCs, where transduction efficiencies approached completeness, while electroporation yielded lower and more variable editing rates (19).

Building on the validated functionality of Cre VLPs in samples from a dual-fluorescent reporter pig (25), we achieved robust activation of latent oncogenic mutations via efficient excision of *LSL* cassettes in organoids derived from oncopigs (28, 29). For these studies, we employed a porcine model of familial adenomatous polyposis and colorectal cancer (CRC) (33), that additionally harbors silenced *KRAS*^*G12D*^ and *TP53*^*R167H*^ mutations and established organoids from polyp biopsy material. Cre-mediated recombination consistently induced a phenotypic transition from branched, differentiated architectures to cystic, highly proliferative morphologies that were maintained over extended culture periods. Notably, comparable structural differences were also observed between human patient-derived organoids from healthy and tumor tissue, respectively (34). We hypothesize that activation of *KRAS*^*G12D*^ and *TP53*^*R167H*^ - orthologous to key driver mutations in human CRC - triggers oncogenic signaling cascades and downstream transcriptional reprogramming, ultimately promoting enhanced proliferation. Future studies should delineate the affected pathways in greater detail and assess autologous reimplantation of edited organoids to accelerate *in vivo* disease modeling (14).

In avian systems, Cre VLPs enabled rapid genome engineering in both chicken *Igh*^*KO*^ DT40 cells and TOCs. This approach provides a practical mean to evaluate disease-resistance genes *ex vivo* (35), complementing earlier mouse-based demonstrations of VLP utility in organ cultures (23) and offering particular value for species in which stable genetic modification remains technically challenging.

Cas9 VLPs enabled robust SKOs in pKDNFs and organoids. Of the two multiplexing strategies evaluated, combining separate single gRNA VLP batches (MPS2) maintained high editing efficiencies across multiple loci and supported complete fragment deletions when flanking gRNAs were applied. In contrast, packaging two gRNAs into a single VLP batch (MPS1) reduced overall editing efficiency and introduced biased gRNA usage. We hypothesize that co-expression of two gRNAs in producer cells leads to competition for Cas9 binding during ribonucleoprotein (RNP) assembly. Differences in gRNA stability or affinity for Cas9 may result in preferential formation of one RNP species (36), which is subsequently enriched during VLP packaging. In addition, the limited cargo capacity of VLPs may further contribute to the observed reduction in editing efficiency, as distinct RNP complexes compete for incorporation into particles. Consequently, MPS2 facilitates the generation of combinatorial genotypes required for modeling CRC consensus molecular subtypes (37-39) and other multigenic disease states in livestock.

Microinjection of multiplexed Cas9 VLPs into the perivitelline space of porcine oocytes yielded a high proportion of efficiently edited blastocysts, thereby simplifying the generation of new genetic lines. In comparison to conventional pronuclear injection - which is technically challenging in porcine oocytes (17) - VLP delivery circumvents the need for pronuclear targeting through biological transduction and mitigates risks associated with vector-encoded nucleases, including off-target editing and mosaicism (40). Relative to our previous workflows employing DNA vectors or RNP complexes (41), Cas9 VLP microinjection increased the proportion of edited blastocysts while reducing mosaicism, underscoring its practical advantages for establishing genetically engineered livestock lines. Notably, no toxicity was observed in porcine cells, organoids, or oocytes, supporting further exploration of direct *in vivo* administration, pending appropriate safety evaluation of the immunogenic potential of VLPs.

In chicken cells, *B2M* targeting with Cas9 VLPs induced a rapid reduction in MFI and thereby in MHC-I expression in HD11 cells within 24 h to 48 h. *In ovo* injection at ED12 resulted in reduced MHC-I expression in the bursa and peripheral blood mononuclear cells (PBMCs). Remarkably, all analyzed bursi (100%) exhibited detectable gene editing, demonstrating the robustness of this approach *in ovo*. Collectively, these findings establish VLP-mediated delivery as a flexible platform for investigating immune and developmental gene function in avian systems, complementing prior *in ovo* VLP vaccine applications (42) and extending the utility of VLPs beyond mammalian models. Notably, the high transduction efficiency (>98%) achieved in PGCs using sfGFP-VLPs enables broad opportunities for the generation of new GM chicken lines. Additionally, gene editing efficiency in PGCs reached approximately 20% 72 h post transduction, indicating sustained Cas9 activity and selective persistence of edited cells in this population.

Multiplexed dCas9 VLPs supported a targeted passive demethylation strategy (30) at the internal *TP53* promoter P2 in porcine cells, achieving greater than 20% reduction at CpG sites using overlapping gRNAs over a six-day period. Delivery of a single gRNA resulted in markedly weaker effects, underscoring the advantage of multiplexing for efficient epigenome editing. Follow-up studies will assess the consequences of targeted demethylation on expression of relevant *TP53* isoforms. In a porcine osteosarcoma model, Niu and colleagues reported that methylation levels at CpGs 2 and 3 within P2 differ by approximately 30-40% between osteoblasts derived from healthy and diseased animals, correlating with increased expression of an oncogenic *TP53* isoform driven by this promoter (43). Based on these findings, the *in vitro* demethylation levels (>20%) achieved in our system are expected to induce a more moderate, yet biologically relevant activation of isoform expression, thereby providing a suitable platform to investigate its functional consequences. Collectively, these findings demonstrate that dCas9 VLP–mediated demethylation extends the application of CRISPR technologies in livestock beyond genome editing towards targeted epigenetic modulation.

To benchmark particle inputs, Cre content in VLP preparations was quantified by ELISA and showed a positive correlation with functional activity. In contrast, Cas9-mediated editing was highly dependent on sgRNA design and target context, necessitating functional pre-testing. Accordingly, we report VLP volumes that consistently performed in pigs and chickens rather than absolute protein loads across all preparations.

While VSV-G confers broad tropism through interaction with the low-density lipoprotein receptor (LDLR), which is widely expressed on the surface of most cell types (44, 45), pseudotyping with the BaEV glycoprotein resulted in reduced efficiency in pigs compared with mice and humans (46), highlighting the importance of species-specific optimization. BaEV utilizes the neutral amino acid transporter ASCT1 as a receptor in human and murine cells. Although this receptor is highly conserved across mammalian species (47), differences in receptor expression levels or compatibility may exist between murine, human, and porcine cells, potentially limiting viral entry in the latter. Future studies should therefore explore alternative envelope glycoproteins to achieve improved tissue or cell-type specificity in pigs and chickens (45).

Finally, although constitutive Cas9-or Cre-expressing livestock lines represent valuable resources (48-50), their generation requires extensive breeding and frequently results in unwanted genotypes. In contrast, direct, time-controlled, and transient delivery of Cre and Cas9 via VLPs can substantially reduce animal use and more closely align with the 3R principles (23). This strategy enables inducible or stage-specific genetic perturbations throughout the animal’s lifespan (51), thereby supporting more refined and flexible experimental designs.

In summary, our results establish VLP-mediated delivery as a safe, rapid, and effective platform for genome and epigenome manipulation across vertebrate classes, including pigs and chickens. By enabling transient and multiplexed delivery of Cre, Cas9, and dCas9 in diverse experimental settings, this approach reduces reliance on stable transgenic lines while substantially expanding experimental flexibility. Collectively, these findings position VLP-based delivery as a practical tool to accelerate functional genetic studies in livestock, with broad relevance for both agricultural and biomedical research.

## Materials and Methods

### Plasmids

For VLP production, pCMV-MMLVgag-3xNES-Cas9 (addgene #181752), pBS-CMV-gagpol (addgene #35614), GAG-CRErec (addgene #119971), CMV-VSV-G (addgene #8454) and SUPERBLADE5 (addgene #134913) were obtained from Addgene (Watertown, USA). The plasmid encoding the BaEV glycoprotein was a kind gift from Emiliano Ricci (CNRS/INSERM, Laboratory of Biology and Modelling of the Cell, École Normale Supérieure de Lyon, France).

### Cloning

The following plasmids were designed and cloned for this study: Gag-dCas9, Gag-sfGFP, and Gag-mCherry. The backbone for Gag-dCas9 was pCMV-MMLVgag-3xNES-Cas9 (addgene #181752). The mutations D10A and H840A disable Cas9’s cutting ability (52), and they were introduced into the plasmid using NEBuilder HiFi DNA assembly. Primer design was done using NEBaseChanger (https://nebasechanger.neb.com/) and the cloning was carried out using the manufacturer’s instructions. Gag-sfGFP and Gag-mCherry were cloned using BIC-Gag-CAS9 (addgene #119942) as a backbone, and Cas9 was excised and replaced with sfGFP from pcDNA3.1_Gag-PCP-sfGFP and mCherry from pcDNA3.1_Gag-N22p_mCherry, respectively. The inserts were amplified using Phusion Polymerase and cloned into the linearized backbone using NEBuilder HiFi DNA Assembly. In any case, correct cloning was checked by Sanger sequencing of the plasmid (Eurofins Genomics, Ebersberg, GER).

### sgRNA design and sequences

The pig and chicken *B2M* sgRNAs have been described previously (48). All other pig sgRNAs were designed using CRISPOR (https://crispor.gi.ucsc.edu/). All sgRNAs were cloned into SUPERBLADE5 using restriction-based cloning as described before (22).

Pig *KDM6A*: 5’ GAAACCTCACGAACCCGAAG 3’

Pig *BRCA2*: 5’ GGTCTCTCTTTGCATCCAAT 3’

Pig *TP53*: 5’ GGCAAAACAGCTTATTGA 3’

Pig *P16*: 5’ GAGGCTAGCCAGTCGGCCGA 3’

Pig *PTEN*: 5’ AGATCGTTAGCAGAAACAAA 3’

Pig *SMAD4*: 5’ ACTATGTACAATGCTCAGAC 3’

Pig *IL10 gRNA2*: 5’ GCTGTTCTCAGACTTAATGC 3’

Pig *IL10 gRNA3*: 5’ TCGGAGTTCCCGGAGCATG 3’

Pig *TP53* demethylation gRNA1: 5’ TGCAGGAGGCGCTGGCACTC 3’

Pig *TP53* demethylation gRNA2: 5’ GCGGGCGCTGTTGCTGCAGG 3’

### Production of VLPs

Protocols for production of Cre and Cas9 VLPs were adapted from recent publications (22, 23). Briefly, 4×10^6^ HEK293-FT cells were seeded into a Corning 10 cm dish (Sigma, Darmstadt, GER) and were transfected on the following day using Lipofectamine 2000 transfection reagent (Thermo Fisher Scientific, Darmstadt, GER) or Viafect (Promega, Madison, USA) according to the manufacturer’s protocol. For producing Cre VLPs, a mixture of Gag-Pol (8.1 µg), VSV-G (3 µg) and Gag-Cre (5.1 µg) was transfected. For producing Cas9 VLPs or dCas9 VLPs, a mixture of Gag-Pol (8.1 µg), VSV-G (3 µg), Gag-Cas9 or Gag-dCas9 (5.1 µg) and sgRNA-SUPERBLADE (13.2 µg) was transfected. For producing sfGFP-/mCherry VLPs, a mixture of Gag-Pol (8.1 µg), VSV-G (3 µg) and Gag-sfGFP or Gag-mCherry (5.1 µg) was transfected. Visible syncytia formation indicated successful co-transfection of all plasmids 24 h to 48 h after transfection (supplementary figure S12). 48 h and 72 h after transfection, VLPs were harvested by removing the supernatant and were stored at 4°C. To purify and concentrate the VLPs, the supernatant was centrifuged at 5,500 xg for 12 h at 4°C or ultracentrifuged at 50,000 xg for 2 h 15 min at 4°C and the resulting VLP pellet was resuspended in cold PBS. The purified VLPs can be stored short term at 4°C and long term at -20°C or -80°C.

### VLP protein content quantification using ELISA

To quantify the protein content in the VLPs, ELISA was performed as previously described (27). For quantification of the Cre amount in VLPs, the following antibodies were used: primary antibody anti-Cre recombinase (D7L7L) XP Rabbit mAb #15036 (Cell Signaling, USA) at a concentration of 19 ng/mL, secondary antibody Goat anti rabbit IgG-HRP #4030-05 (SouthernBiotech, Birmingham, USA) at a concentration of 23.75 ng/mL. For quantifying the Cas9 amount in VLPs, the following antibodies were used: primary antibody mouse monoclonal IgGiK anti-Cas9 AB (7A9-3A3) (Cell Signaling, USA) at a concentration of 0.4 µg/mL, secondary antibody mouse igGk BP-HRP sc-516102 (SantaCruz, Dallas, USA) at a concentration of 40 ng/mL. Three batches of distinctly produced Cre and Cas9 VLPs were measured and tested for functionality. For consistency of the experiment, typically 5-10 µL Cre VLPs and 10-20 µL Cas9 VLPs were used. It should be noted that the efficiency of all VLPs varies depending on the transduction of pig or chicken cells, which is why each batch was tested on the respective target cells to ensure maximal reliability and performance.

### Transduction of porcine cells

To test VLPs on pKDNFs, 150,000 cells were seeded in a 12-well plate. On the following day, medium was changed and 5-20 µL VLPs were added. Analysis was conducted starting 24 h post transduction at distinct time points.

For Cre VLPs, pKDNFs from a dual fluorescent reporter pig were used (25), which switch from mTomato to eGFP expression upon Cre-mediated recombination. Cells were transduced with 10 µL Cre VLPs and fluorescence microscopy (ECHO Revolve Version v7.2.2, San Diego, USA) was conducted after 24 h, 48 h, and 72 h.

To test Cas9 VLPs, 20 µL were applied to the cells. 48 h post transduction, cells were detached and gDNA isolated using QuickExtract™ DNA Extraction Solution (Biosearch Technologies, Petaluma, USA). PCR amplified a 200 – 700 bp region around the target locus (sequences in supplementary table S1). The product was checked for correct size on an agarose gel and subsequently sent for Sanger sequencing (Eurofins Genomics, Ebersberg, GER). Sequencing results were analyzed for the percentage of INDELs using the online ICE analysis tool by Synthego (https://ice.editco.bio).

To test reporter VLPs, 5 µL sfGFP-VLPs or 10 µL mCherry VLPs were added to the cells. sfGFP or mCherry expression was analyzed 24 h and 48 h after transduction using fluorescence microscopy (ECHO Revolve Version v7.2.2, San Diego, USA) and Flow cytometry (Attunes NxT, Thermo Fisher Scientific, Darmstadt, GER).

For demethylation, pKDNFs with high methylation levels at the target locus were used. 300,000 cells were seeded in a 6-well plate and transduced every second day with 20 µL of dCas9 VLPs for up to 15 days. Cells were harvested after 1, 3, 6, and 15 days for methylation analysis as described below, with remaining cells reseeded for continued treatment.

### Transduction of avian cells

To test Reporter VLPs, 1×10^6^ DT40 cells were seeded in a 12-well plate. The next day, 5 µL of sfGFP VLPs or 10 µL of mCherry VLPs were added to the medium. For PGCs, 125,000 cells were seeded in a 12-well plate and 5 µL, 10 µL, or 20 µL sfGFP VLPs were applied. Fluorescent protein expression was analyzed 24 h and 48 h after transduction using fluorescence microscopy (ECHO Revolve Version v7.2.3, San Diego, USA) and Flow cytometry (Attunes NxT, Thermo Fisher Scientific, Darmstadt, GER).

Chicken *Igh*^*KO*^ DT40 cells (26) were used to test Cre VLPs in chicken suspension cells, as they exhibit an eGFP cassette flanked by loxP sites. 65,000 cells were seeded in a 12-well plate and on the next day 5 µL, 10 µL, and 20 µL Cre VLPs were added. eGFP expression was analyzed 24 h, 48 h, and 72 h after transduction using Flow cytometry (Attunes NxT, Thermo Fisher Scientific, Darmstadt, GER) and fluorescence microscopy (Discover ECHO Revolve v7.2.3, San Diego, USA).

To evaluate *B2M* Cas9 VLPs, 250,000 adherent chicken HD11 cells (53) were seeded in 12-well plates and transduced the following day with 10 µL Cas9 VLPs. Cells were harvested 48 h and 72 h post transduction using PBS/EDTA and stained for flow cytometry with mouse anti-chicken MHC-I primary antibody (2.5 µg/mL) and IgG-APC secondary antibody (2 µg/mL). Data were acquired on an Attune NxT flow cytometer (Thermo Fisher Scientific, Darmstadt, Germany). For PGCs, 125,000 cells were seeded per well in 12-well plates, and cell pellets were collected after 48 h and 72 h to assess gene editing. gDNA from HD11 cells and PGCs was isolated using the ReliaPrep Blood gDNA Miniprep System (Promega, Madison, USA) according to the manufacturer’s instructions. Gene editing was verified by PCR using FIREPol DNA Polymerase Mastermix (Solis Biodyne) with an annealing temperature of 60°C and a 30 s elongation time; chicken *B2M* primers are shown in supplementary table S1. PCR products were size-verified by agarose gel electrophoresis and subjected to Sanger sequencing (Eurofins Genomics, Ebersberg, Germany). INDEL frequencies were quantified using the TIDE web tool (https://tide.nki.nl).

### Porcine organoid culture and transduction of organoids

Permission for taking biopsies from pigs was issued by the government of Upper Bavaria, Germany (ROB-55.2-2532.Vet_02-23-82). Experiments were performed according to the German Welfare Act and European Union Normative for Care and Use of Experimental Animals. Porcine colonic organoids were established from a biopsy of either normal mucosa or a polyp from a pigs’ colon as described before (48) with adaptations to the protocol. Briefly, the tissue was minced and washed five times with PBS containing 1% Penicillin/Streptomycin. The tissue pieces were digested using preheated 10 mg/mL collagenase P (Sigma, Darmstadt GER) for 1 h at 37°C. Afterwards, the tissue was resuspended in cold AIM V (Thermo Fisher Scientific, Darmstadt, GER) and the crypts were released mechanically by shaking for 30 sec. The suspension was passed through a 100 µm cell strainer to remove unwanted residues. The crypts were pelleted for 5 min at 400 xg and plated in a prewarmed 24 well plate with Matrigel (Corning, New York, USA) with a density of 200 - 400 crypts per well. After 20 min at 37°C, organoid growth medium (human) (OGM, Stemcell Technologies, Vancouver, CAN) with 10 mM Thiazovivin (Sigma, Darmstadt, GER) and 100 µg/mL Primocin (Invitrogen, Thermo Fisher Scientific, Darmstadt, GER) was added. After 10 days in culture, Thiazovivin and Primocin were removed from the medium. To passage organoids, the medium was aspired, and the domes dissolved with 1 mL cold PBS. The samples were pelleted and resuspended in cold PBS. The organoids were dissociated using a 25 G syringe and incubated with gentle dissociation reagent (Stemcell Technologies, Vancouver, CAN). The cells were again centrifuged and plated as described above.

Protocols for transduction of organoids with VLPs were adapted from previous publications (27). Briefly, organoids were dissociated and centrifuged for 5 min at 400 xg. The pellet was resuspended in 500 µL OGM with Thiazovivin. The suspension was distributed to a 96-well plate with 100 µL per well and VLPs were added. The plate was then centrifuged at 200 xg and room temperature for 3 h and then incubated for 3 h at 37°C. Finally, the suspensions were mixed, briefly centrifuged and plated again.

To test Cre VLPs, porcine colonic organoids were transduced with 5 µL of Cre VLPs. In case of a fluorescence readout, eGFP expression was analyzed 24 h, 48 h and 72 h after transduction using fluorescence microscopy (ECHO Revolve Version v7.2.2, San Diego, USA). In case of activation of latent oncogenic mutations, DNA and RNA were extracted from the organoids using the Qiagen Mini Kit following the manufacturer’s instructions. One dome of organoids was dissociated, centrifuged and resuspended in 350 µL lysis buffer. Concentrations of DNA and RNA were measured, and the samples stored immediately at -20°C and -80°C, respectively. DNA was submitted to recombination PCR and RNA to pyrosequencing, which is described below. Furthermore, the morphology of the organoids was assessed until 46 days after transduction.

To test Cas9 VLPs, porcine organoids were transduced with 20 µL of Cas9 VLPs. After 5 days, DNA was extracted and analyzed for INDELS as described above.

### Methylation analysis

pKDNFs with high methylation levels in the desired promoter 2 (P2) region of *TP53* were transduced with dCas9 VLPs. gDNA was extracted prior to and after transduction using QuickExtract™ DNA Extraction Solution (Biosearch Technologies, Petaluma, USA) and concentration was measured using FastGene NanoSpec Photometer (Nippon Genetics, Dueren, GER). For bisulfite conversion, the EZ DNA Methylation-Direct™ Kit (Zymo Research, Irvine, USA) was used according to the manufacturer’s protocol starting from step 2. 400 ng gDNA were converted per sample. The reagents from the PyroMark PCR kit (Qiagen, Hilden, GER) were used to amplify the target region using 3 µL of CT-converted DNA as input. After confirmation of successful PCR, pyrosequencing was conducted using PyroMark Q48 Advanced CpG reagents on a PyroMark Q48 Autoprep instrument (Qiagen, Hilden, GER). The *TP53* CpG Methylation assay was designed using PyroMark Assay Design 2.0 software (Qiagen, Hilden, GER).

### Recombination PCR

To verify the activity of Cre delivered by VLPs, a PCR across the *LSL* cassette in intron one of both *KRAS* and *TP53* was conducted. Isolated gDNA of wildtype and VLP-transduced organoids was used. GoTaq® G2 DNA Polymerase (Promega, Walldorf, GER) was used for amplification and the following cycle conditions were set: 3 min 95°C, then 40 cycles of 45 sec, 95°C, 45 sec, 59°C, 90 sec, 72°C followed by 5 min at 72°C.

### Pyrosequencing

To prove that the latent mutations were activated and expressed by VLP delivered Cre, the proportion of wildtype (G) and mutant (A) *KRAS* and *TP53* alleles was assessed. Therefore, RNA of wildtype and VLP-transduced organoids was isolated, and cDNA was synthesized using the LunaScript RT Master Mix Kit (NEB) according to the manufacturer’s protocol. cDNA was diluted (1:3) and the desired loci amplified with the PyroMark PCR kit (Quiagen, Hilden, GER). PCR fragments were checked on an agarose gel before submitting them to pyrosequencing. *KRAS* and *TP53* allelic quantification assays were designed using PyroMark Assay Design 2.0 software (Quiagen, Hilden, GER) and pyrosequencing was conducted using the PyroMark Q48 Advanced CpG reagents on a PyroMark Q48 Autoprep instrument (Quiagen, Hilden, GER).

### *In vitro* embryo culture and microinjection of VLPs into porcine oocytes

Oocytes were prepared and submitted to microinjection as described before (17, 54). Briefly, oocytes were isolated from prepubertal gilts (Vion SBL Food GmbH, Landshut, Germany) and maturated *in vitro*. The zygotes were either *in vitro* fertilized or submitted to parthenogenesis. In the first case, successfully maturated zygotes were determined visually, and deep-frozen sperm was thawed, prepared, and assessed for viability. Sperm was transferred to the zygotes and incubated in a triple-gas incubator at 38.5°C for 7 h. Afterwards, the zygotes were washed and selected for the presence of two polar bodies indicating successful fertilization. The fertilized embryos were submitted to microinjection, where approximately 10 pL of VLPs were injected into the perivitelline space (22). Embryos were cultivated for 7 days until blastocysts developed for the assessment of targeting efficiency. For parthenogenesis, following microinjection oocytes with polar bodies were submitted to activation by applying a 43 V DC current for 80 µs. Fused oocytes were washed and incubated with 5 µg/mL Cytochalasin B to prevent extrusion of the second polar body. After 3 h, activated oocytes were washed and cultivated for 7 days.

To assess the impact on embryo development, different dilutions of VLPs were injected into zygotes prior to parthenogenesis (supplementary table S2). Subsequently, VLPs were injected into zygotes that were *in vitro* fertilized (supplementary table S3). To analyze gene editing efficiencies, blastocysts were transferred into a PCR tube containing 12 µL embryo lysis buffer and incubated at 65°C for 1 h followed by 95°C for 10 min. Subsequently, the lysate was used as template for ICE analysis as described above.

### Transduction of TOCs

TOCs were prepared as described before (35) from embryonated JH-KO chicken eggs (9) (TUM Animal Research Center, Freising, GER) that exhibit an eGFP cassette flanked by loxP sites. According to European regulations, interventions and treatments involving chicken embryos are not classified as animal experiments and are even recognized as a replacement method under the 3Rs principle. In accordance with Article 14 of the German Animal Welfare Ordinance (TierSchVersV), ethical approval is only required if chicken embryos are used at a developmental stage before hatching and are intended to survive beyond hatching. Therefore, terminal procedures such as organ harvesting prior to hatching do not require ethical review or authorization as animal experiments. Briefly, eggs were embryonated until ED20, embryos were decapitated, and the tracheae were removed. The tracheae were cut into 0.8 mm rings and transferred into a falcon holding 300 µL TOC Medium (Medium 199 with Hanks’ salts, 1% Penicillin/Streptomycin, 1% Glutamaxx). 10 µL Cre VLPs were added into a falcon containing n=3 TOCs in medium. eGFP expression was analyzed 48 h after transduction using fluorescence microscopy (Discover ECHO Revolve Version v7.2.2, San Diego, USA).

### *In ovo* injection of VLPs

Wild type eggs (White Leghorn, TUM Animal Research Center, Freising, GER) were collected and stored up to 3 weeks at 16°C. For *in ovo* experiments, eggs were incubated at 37.8°C, after 7 days the eggs were candled and infertile or dead eggs were removed. 50 µL VLPs were injected into a blood vessel on ED 12 as described before (55). On ED 19 blood was collected again *in ovo*, subsequently the embryo was decapitated and bursi were collected. PBMCs were isolated from blood using Histopaque (Sigma, Darmstadt, GER) and consequently were stained for flow cytometry analysis. Bursi were strained through a cell strainer, density centrifugation using Histopaque (Sigma, Darmstadt, GER) was performed and subsequently resulting cells were stained for flow cytometry as described before. gDNA was isolated from bursa cells using ReliaPrepTM Blood gDNA Miniprep System Kit (Promega. Madison, USA) according to the manufacturer’s protocol. gDNA was analyzed for CRISPR edits as described above.

## Funding

This project was funded by the Deutsche Forschungsgemeinschaft (DFG, German Research Foundation) in the framework of Collaborative Research Centre Microbiome Signatures (SFB1371), project number 395357507 (awarded to TF), and the Research Unit ImmunoChick (FOR5130), project number 434524639 (awarded to BS and TvH).

## Acknowledgments

We would like to thank Jeff Moore, Jay Moore and Yiting Wu (Curileum) for providing us with a protocol for improved organoid establishment from colon biopsies. ChatGPT (https://chat.openai.com/) was used to improve conciseness of language and coding, with no contribution to the scientific content. BioRender was used to generate illustrations (https://BioRender.com).

## Notes

**Competing Interest Statement:** The authors declare that the research was conducted in the absence of any commercial or financial relationships that could be construed as a potential conflict of interest.

### Competing Interest Statement

The authors have declared no competing interest.

