## Supporting Information for "Virus-Like Particles: The Next Frontier in Livestock Gene Editing"

**This PDF file includes:**

Figures S1 to S12
Tables S1 to S3

**Figures**

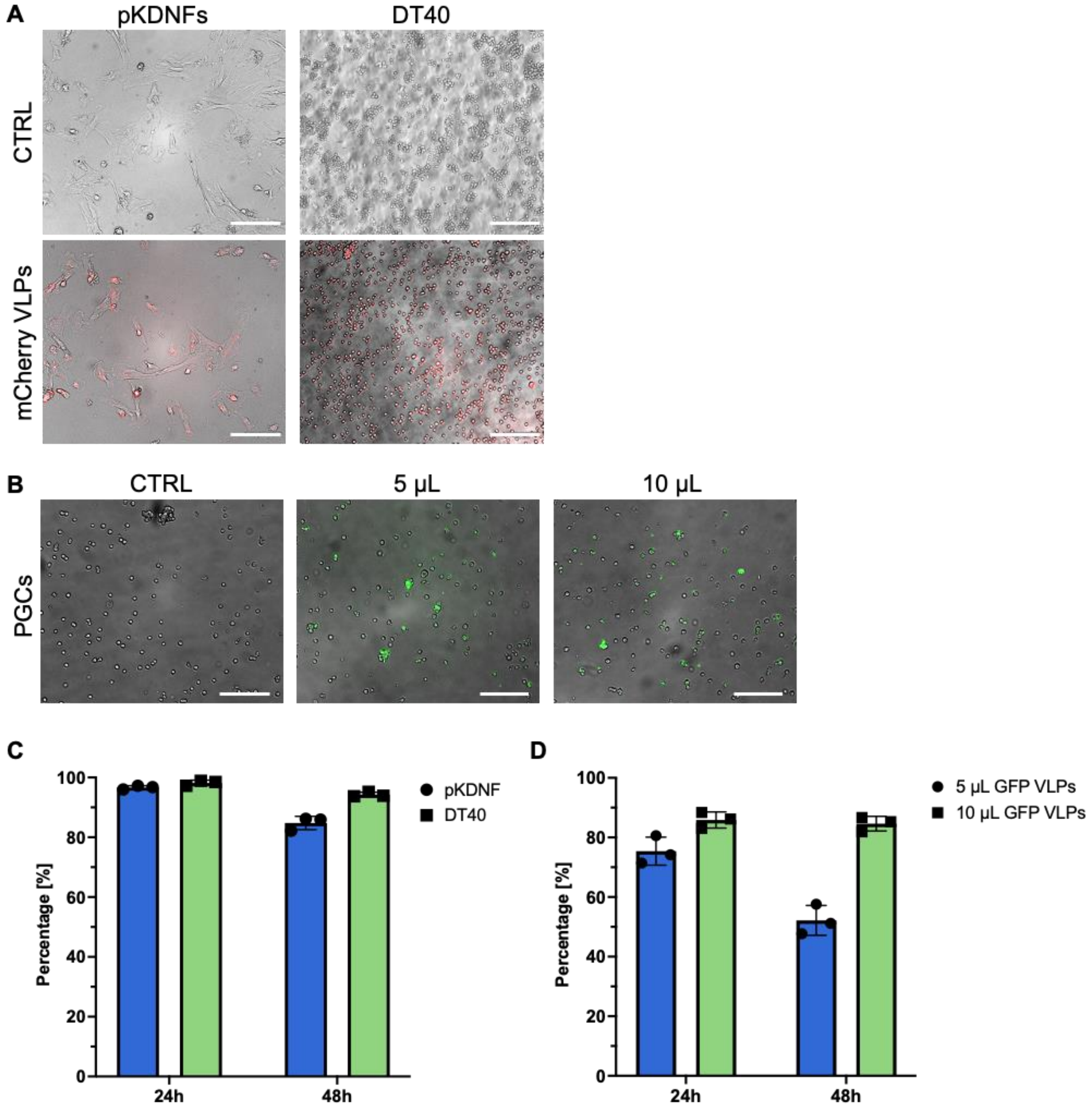


**Figure S1. VLPs efficiently package and transport fluorescence proteins and release them into various target cells. A**: Representative fluorescence microscopy images of porcine pKDNFs and DT40 chicken cells transduced with 10 µL mCherry VLPs 24 h post transduction. Scale bar = 130 µm. **B**: Representative fluorescence microscopy images of PGCs transduced with different volumes of sfGFP VLPs 24 h post transduction. Scale bar = 130 µm. **C**: Relative count of fluorescence expression in single cells from DT40 cells and pKDNFs transduced with 10 µL mCherry VLPs 24 h and 48 h post transduction compared to control (n=3; SD shown). **D**: Relative count of fluorescence expression in single cells from PGCs transduced with 5 µL and 10 µL sfGFP VLPs 24 h and 48 h post transduction compared to control (n=3; SD shown).

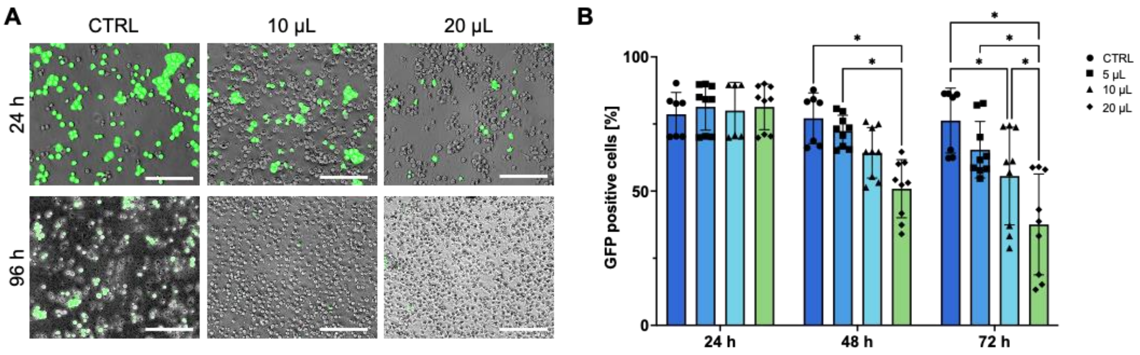


**Figure S2. VLPs efficiently deliver functional Cre to chicken *in vitro* and *ex vivo* culture systems. A**: Representative fluorescence microscopy pictures of *Igh*$^{KO}$ DT40 chicken cells transduced with Cre VLPs in different volumes (10 µL and 20 µL) at 96 h post transduction. Scale bar = 230 µm. **B**: eGFP expression of *Igh*$^{KO}$ DT40 cells transduced with Cre VLPs in different volumes (5 µL, 10 µL, and 20 µL) at 24 h, 48 h, and 72 h post transduction (n=9; p<0,05; Multiple ANOVA; SD shown).

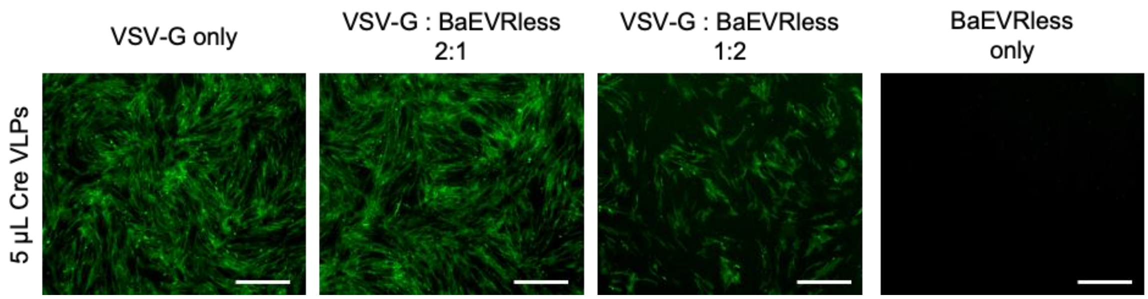


**Figure S3. Pseudotyping VLPs with a baboon envelope glycoprotein reduces efficiency of transduction in porcine cells.** Cre VLPs were pseudotyped with different ratios of VSV-G and a BaEV, which is reported to enhance transduction efficiency in murine cells. In porcine cells, an adverse effect is observed. Fluorescence images taken 48 h post transduction. One representative experiment shown, n > 3. Scale bar = 430 µm.

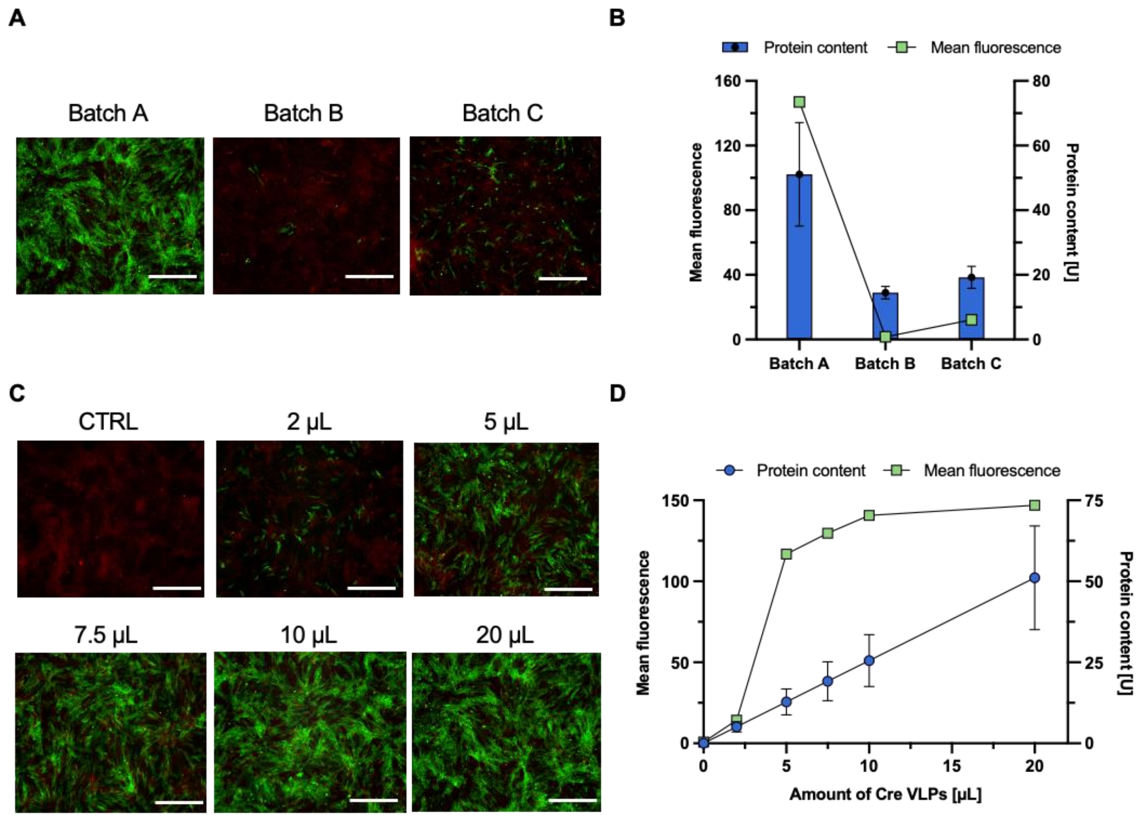


**Figure S4. Results of Cre VLP functionality tests and corresponding protein quantities. A** and **B**: 20 µL of three distinctly produced batches of Cre VLPs were submitted to functional tests on reporter pKDNFs (**A**); the fluorescence of eGFP was analyzed using ImageJ and the VLP batches were submitted to ELISA to quantify the amount of Cre protein in the VLPs (**B**). **C** and **D**: Cre VLP Batch A was used to determine the minimal amount of Cre VLPs needed to efficiently excise loxP sites. Different amounts were tested on reporter pKDNFs (**C**); the fluorescence of eGFP evaluated using ImageJ and the results plotted against the corresponding amount of Cre protein (**D**). All images taken 72 hours post transduction, scale bar = 430 µm.

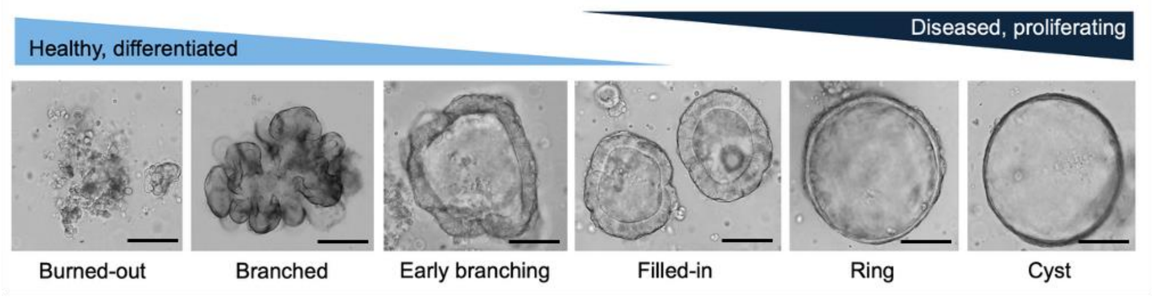


**Figure S5. Morphology of porcine colon organoids differs depending on the disease state of the cells.** Organoids derived from healthy colon tissue show a differentiated, branched phenotype whereas organoids that are derived from diseased tissue appear in a cystic, proliferating shape. Scale bar = 70 µm.

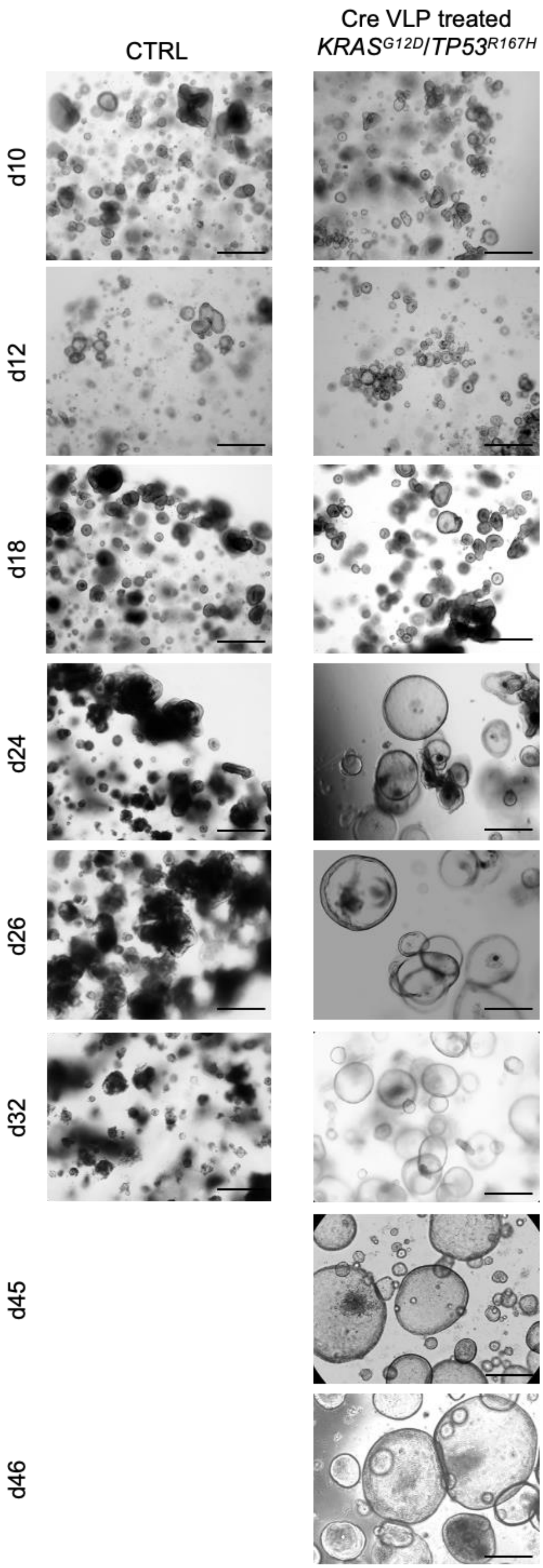
CTRL
Cre VLP treated
$KRAS^{G12D}/TP53^{R167H}$
d10
d12
d18
d24
d26
d32
d45
d46

**Figure S6. Brightfield images of organoids that were transduced with 5 µL Cre VLPs show a change in morphology when grown in culture for >24 days.** Latent oncogenic mutations in *KRAS* and *TP53* were activated. After longer culture periods, a shift in their phenotype was observed, from a branched, more differentiated to a cystic, proliferating morphology. One representative experiment shown, n > 2. Scale bar = 130 µm.

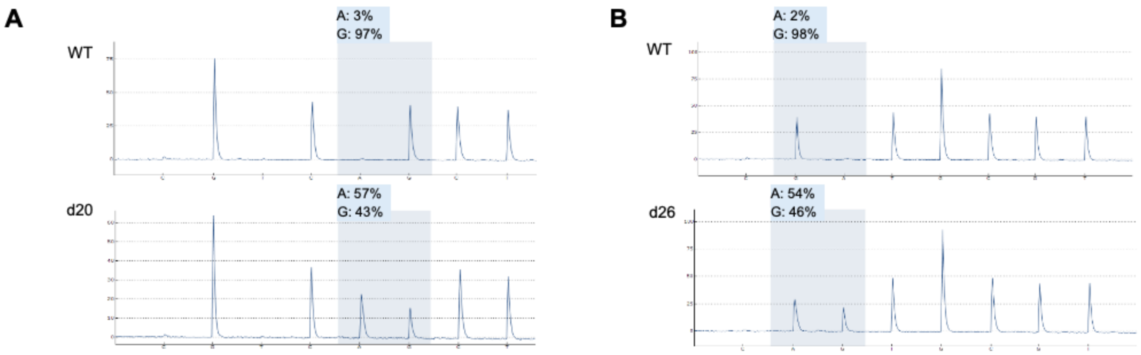


**Figure S7. Allelic quantification reveals activation of latent oncogenic mutations on cDNA level.** Organoids with latent oncogenic mutations were transduced with Cre VLPs and afterwards, cDNA was subjected to pyrosequencing. Allelic quantification of cDNA from *TP53* (**A**) and *KRAS* (**B**) confirms the expression of the mutant allele on RNA level. In the blue box: G represents WT allele, A represents the mutant allele.

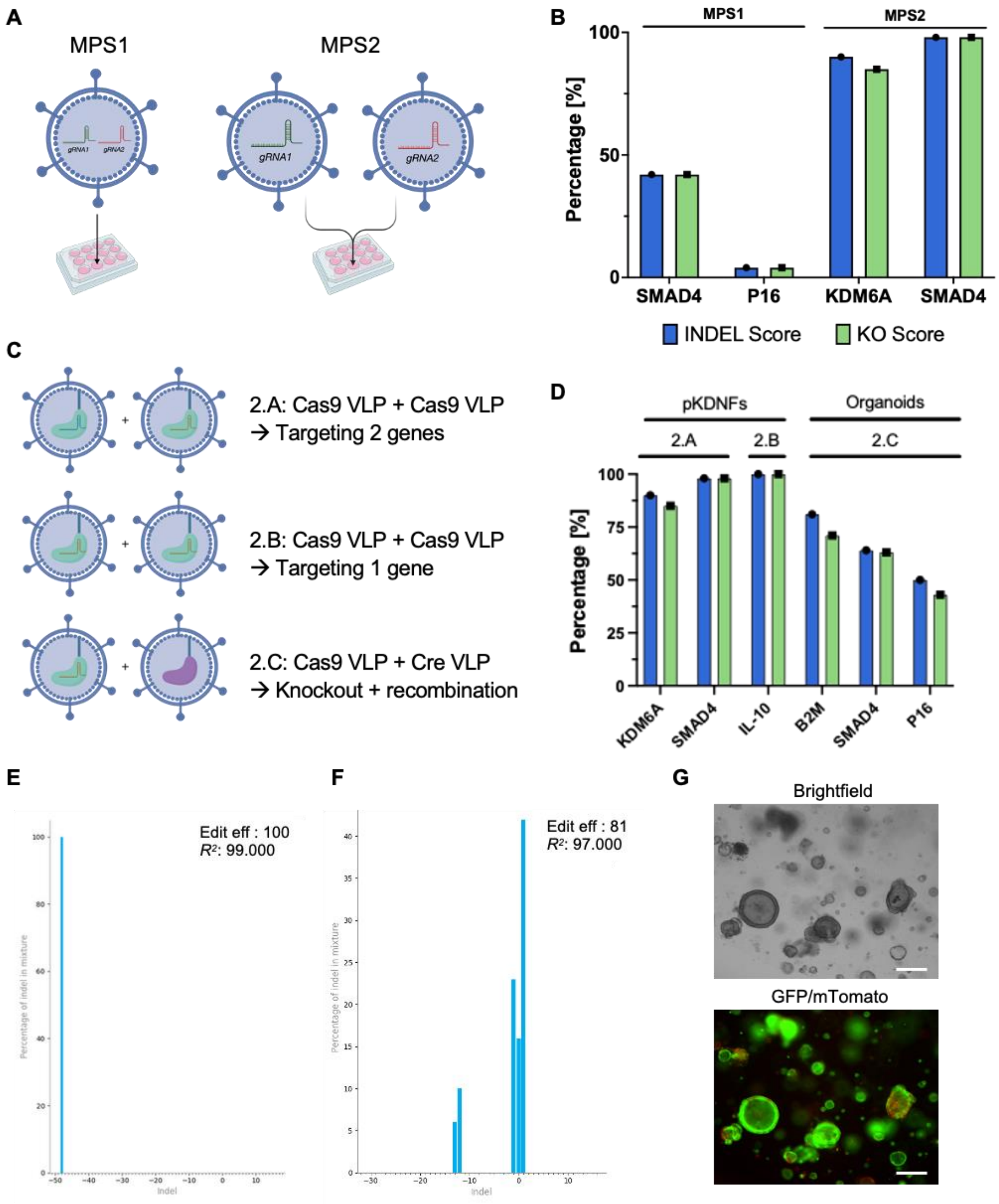


**Figure S8. Results of different Cas9 VLP multiplexing strategies on porcine fibroblasts and organoids. A**: Depiction of the two multiplexing strategies (MPS) tested in this study. Images created using bioRender. **B**: Bar graph showing that MPS1 results in overall loss of editing efficiency as well as a bias of one gRNA over the other. MPS2 leads to high KO scores in both targeted loci, comparable to SKOs. Shown is one representative replicate per gene. Method-validation experiments confirmed reproducible efficiencies across biological and technical replicates. **C**: Schematic overview of MPS2 variations: (2.A) multiplexing two Cas9 VLP batches to target two genes simultaneously; (2.B) multiplexing two Cas9 VLP batches to target one gene with two flanking gRNAs; (2.C) multiplexing Cas9 and Cre VLPs to achieve site-specific recombination and a targeted knockout simultaneously. **D**: Bar graph showing high KO efficiencies of MPS2 in porcine pKDNFs and organoids using different variation of MPS2. One representative replicate per gene is shown; validation experiments confirmed reproducible efficiencies across biological and technical replicates. **E**: ICE analysis for the *IL-10* KO on pKDNFs using MPS2. The INDEL graph shows a complete fragment deletion. **F** and **G**: Exemplary analysis of organoids transduced with multiplexed Cas9 and Cre VLPs, targeting *B2M* and the dual fluorescent reporter cassette. The INDEL graph (**F**) depicts the Cas9-mediated edits and representative fluorescence images at 48 h post transduction (**G**) show Cre-mediated eGFP expression. Scale bar = 130 μm.

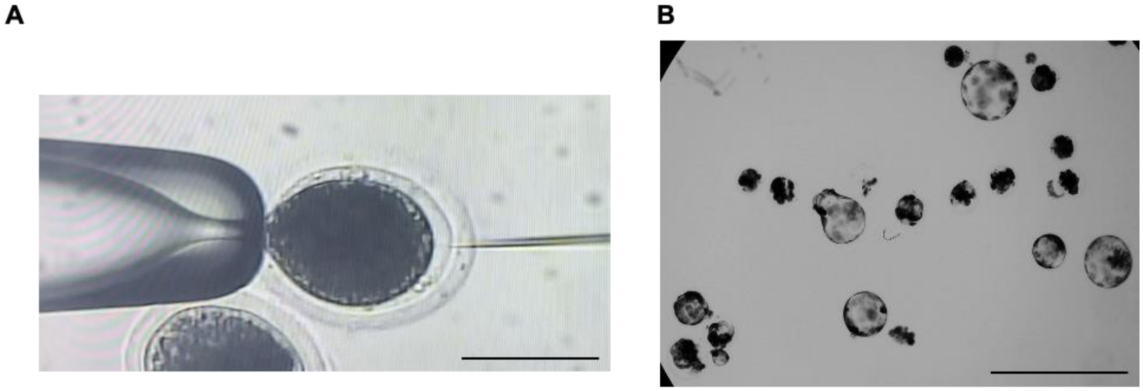


**Figure S9. Microinjection of multiplexed Cas9 VLPs into porcine oocytes and development into blastocysts. A**: The injection needle of the micromanipulator penetrating a porcine oocyte to inject Cas9 VLPs into the perivitelline space. Scale bar = 130 µm. **B**: After 7 days in culture, blastocysts start to develop and can be analyzed for genome editing. Scale bar = 630 µm.

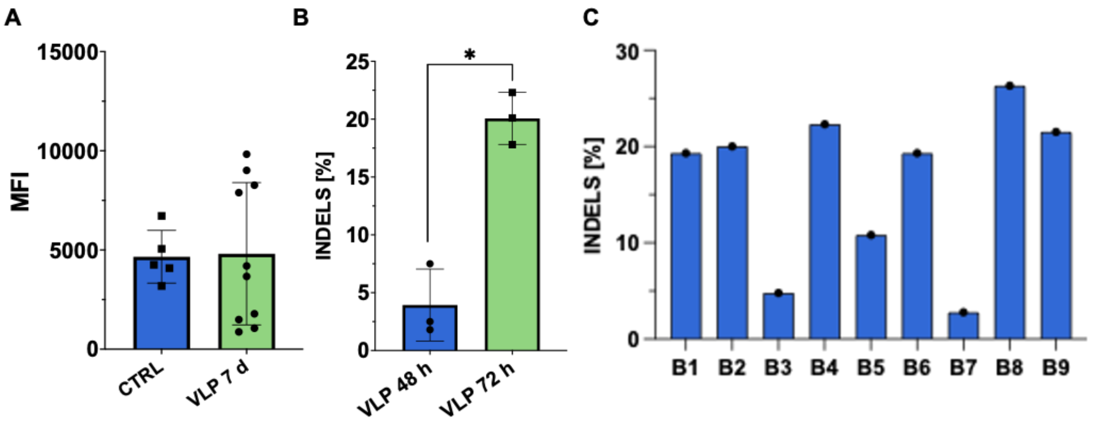


**Figure S10. VLP-mediated gene editing decreases MHC-I expression on the surface of chicken cells *in vitro* and *in ovo*. A**: Mean fluorescence intensity (MFI) of MHC-I expression in PBMCs from ED19 WT chicken embryos transduced with 50 µL Cas9 VLPs targeting *B2M* 7 days post transduction compared to uninjected control (n≥5; p<0,05; Multiple ANOVA; SD shown). **B**: Bar graph showing INDEL scores of Cas9 VLPs targeting *B2M* in chicken PGCs 48 h and 72 h post transduction (n=3; t-test; SD shown). **C**: Bar graph showing INDEL scores from cells from nine individual ED19 bursa from WT chickens transduced with Cas9 VLPs targeting *B2M* 7 days post transduction.

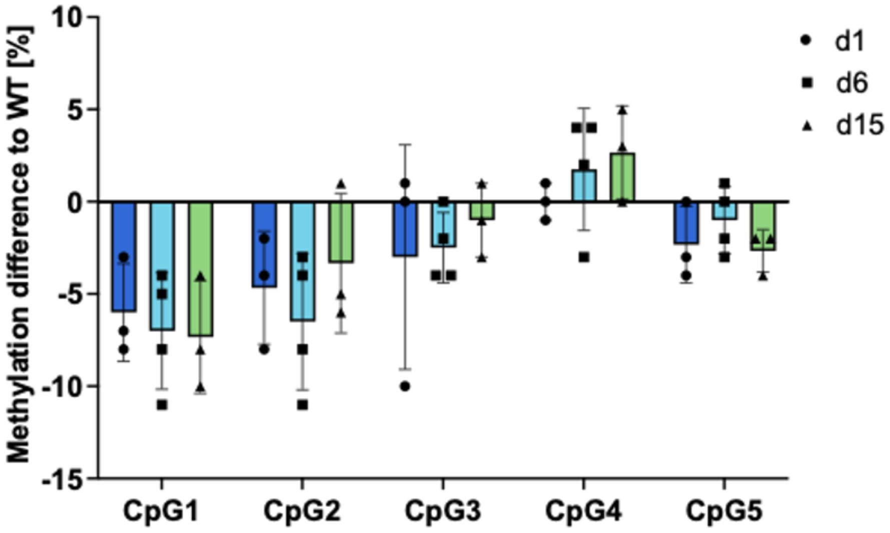


**Figure S11: Demethylation of internal promoter P2 of *TP53* in pKDNFs using only gRNA1 in dCas9 VLPs.** pKDNFs were transduced with 20 µL dCas9 VLPs every second day and analyzed on d1, d6 and d15 post transduction. Graph depicts demethylation difference to WT in CpG1-5 (n >= 3, SD shown).

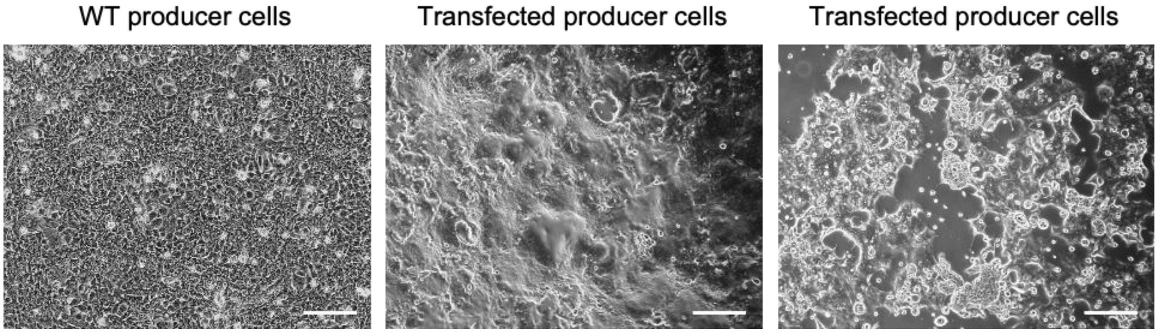


**Figure S12: Syncytia formation indicates successful transfection for VLP production.** Representative microscopic images show untransfected HEK 293 FT cells (left) or HEK 293 FT cells 48 h (middle) and 72 h (right) post transfection. Scale bar = 430 µm.

**Tables**

**Table S1.** All primer sequences used in this study.

| Primer Name | Sequence 5’ to 3’ | Purpose |
|---|---|---|
| Chicken B2M fwd | CAAGGTGCAGGTGTACTCC | TIDE analysis |
| Chicken B2M rev | ACTTGTAGACCTGCGGCTC | TIDE analysis |
| Pig B2M fwd | CCACCCAGTCCAACCTTTGCC | ICE analysis |
| Pig B2M rev | CCAGAGTTAGCGCCCGGAGT | ICE analysis |
| Pig KDM6A fwd | CGCTTTCGGTGATGAGGAAA | ICE analysis |
| Pig KDM6A rev | TGGCTCGAATTTGGAAAAGC | ICE analysis |
| Pig SMAD4 fwd | AGCAATTTCATCTTTTCCCAAGT | ICE analysis |
| Pig SMAD4 rev | GACTAACCTGAAGCCTCCCA | ICE analysis |
| Pig P16 fwd | CTCCGGGTGGAAAGATACCG | ICE analysis |
| Pig P16 rev | TGTCCTCCGACGGATTTCTG | ICE analysis |
| Pig PTEN fwd | GTCGCTGCAACCATCCAG | ICE analysis |
| Pig PTEN rev | AGCCTAGGGTGTGTTCATCT | ICE analysis |
| Pig TP53 KO fwd | ACCCTGGTCCCAAAGTTGAA | ICE analysis |
| Pig TP53 KO rev | TATAGCGATGGTGAGTGGGC | ICE analysis |
| Pig BRCA2 fwd | GCGGAATCTTGCTCTTTTGGA | ICE analysis |
| Pig BRCA2 rev | GCCAGTCTCCTCTTTGTCCA | ICE analysis |
| Pig IL-10 fwd | GGCCTCACTGAACCCACAAT | ICE analysis |
| Pig IL-10 rev | CCAACCACGTCCAACTCTTG | ICE analysis |
| Pig TP53 DM fwd | AGTTAAGAATTGGTTGGATGAAAATTTA | Methylation analysis |
| Pig TP53 DM biot rev | AAACCAATCCCTCAAAACCACTAACC | Methylation analysis |
| Pig TP53 DM pyroseq | GGTTGGATGAAAATTTAGATG | Methylation analysis |
| Pig KRAS rec fwd | AAAGCGGTACTTGCCTTTAAT | Recombination PCR |
| Pig KRAS rec rev | TGAGGAAAAGAACAGTGCAAA | Recombination PCR |
| Pig TP53 int1 fwd | TGAGGAATTTGTATGCCAAGG | Recombination PCR |
| Pig TP53 rec rev | TTCCACCACTGAATCCACAA | Recombination PCR |
| Pig KRAS cDNA fwd | GCCTGCTGAAAATGACTGA | Allelic quantification |
| Pig KRAS cDNA biot rev | CATGTACTGGTCCCTCATT | Allelic quantification |
| Pig KRAS pyroseq | TGTGGTAGTTGGAGCTG | Allelic quantification |
| Pig TP53 cDNA fwd | TGTCCGCGCCATGGCCATCT | Allelic quantification |
| Pig TP53 cDNA biot rev | CAGAGCCGACCTCGGGCG | Allelic quantification |
| Pig TP53 pyroseq | ACCGAGGTGGTGA | Allelic quantification |
| dCas9-mut fwd-1 | CATCGGCCTGGCCATCGGCACCA | dCas9 mutagenesis |
| dCas9-mut rev-1 | CTCTGAGGCACGATGGCGTCCACATCG | dCas9 mutagenesis |

| | | |
|---|---|---|
| dCas9-mut fwd-2 | CGATGTGGACGCCATCGTGCCTCAGAG | dCas9 mutagenesis |
| dCas9-mut rev-2 | TGGTGCCGATGGCCAGGCCGATG | dCas9 mutagenesis |
| dCas9 screening fwd-1 | CTGCAAAGAAAAGGGGCACT | dCas9 screening |
| dCas9 screening rev-1 | CTTCTTGATGCTGTGCCGGT | dCas9 screening |
| dCas9 screening fwd-2 | TCCACGACGACAGCCTGACC | dCas9 screening |
| dCas9 screening rev-2 | GATTTCCTGCTCGCTCTTGG | dCas9 screening |

**Table S2.** Injection of different VLP dilutions into oocytes that were submitted to parthenogenesis to assess VLP toxicity on the development on blastocysts.

| Dilution of VLPs:PBS | 1:2 | 1:4 | 1:50 |
|---|---|---|---|
| # of injected oocytes | 57 | 74 | 69 |
| # of developed blastocysts | 17 | 11 | 17 |
| % of developed blastocysts | 29.825 | 14.865 | 24.638 |

**Table S3.** Injection of undiluted, multiplexed VLPs into zygotes that were *in vitro* fertilized. ICE analysis revealed efficient gene editing.

| Injection type | VLPs |
|---|---|
| # of injected oocytes | 127 |
| # of developed blastocysts | 6 |
| % of developed blastocysts | 4.724 |
| # of edited blastocysts | 4 |
| % of edited blastocysts | 66.667 |
| % INDELs | 96 - 100 |
